# Microscale assay to evaluate the minimum inhibitory concentration of purified compounds with limited sample volume

**DOI:** 10.64898/2026.07.07.737130

**Authors:** Shashank Kashyap, Sumit Biswas

**Author notes:** Corresponding author Email id.

## Abstract

The minimum inhibitory concentration (MIC) is a standard measure for describing the lowest effective dose concentration of an antimicrobial compound in clinical practice; yet, conventional assays often require a substantial amount of antimicrobial compound, limiting their use with scarce, purified agents. Here, we describe a simple and reproducible technique to evaluate the MIC for purified compounds with a limited sample size. The protocol describes the MIC steps against a bacterial strain while minimizing the use of reagents and materials. It is helpful for screening purified natural products as antimicrobial agents and in early-stage drug discovery. The protocol adapts standard microplate-based assays for two-fold dilution of the compound, ensuring their applicability in microbiological studies. The MIC value of the standard antibiotic kanamycin against *Staphylococcus aureus, Vibrio fischeri, Klebsiella pneumoniae*, and *Escherichia coli* was determined using our method, and was found to be consistent with the conventional broth microdilution method, validating its reliability. Therefore, this method offers a practical and viable solution for antimicrobial drug discovery, addressing the disparity between limited compound availability and comprehensive microbiological assessment of MIC.

## Introduction

With the increasing threat of antimicrobial resistance worldwide and the advancement in drug discovery, there is an increasing demand for investigating natural sources for the development of novel antimicrobials and their screening approaches [1, 2]. Currently, many investigations are focused on developing and isolating new antimicrobial compounds or peptides from plants, microorganisms, and other biological organisms. These natural compounds often demonstrate promising antimicrobial activity, prompting further investigations into their potential therapeutic applications [3]. However, determining the Minimum Inhibitory Concentration (MIC) is crucial in evaluating their efficiency, and it is a key pharmacodynamic parameter that quantifies the *in vitro* efficacy of an antimicrobial agent by determining the lowest concentration of a compound to inhibit the growth of specific microorganisms in standardized laboratory conditions [4, 5]. Efficient and reproducible MIC determination is the key factor for several reasons. Primarily, it provides a baseline measure for comparing the activity of different antimicrobial agents against the same organism. Furthermore, it facilitates the establishment of susceptibility breakpoints, which are essential for guiding clinical treatment decisions and monitoring the emergence of resistance. In addition, the MIC serves as a vital parameter for understanding the dose-response relationship and for predicting the concentrations of the antimicrobial agent required to achieve therapeutic efficacy *in vivo* [6, 7].

Despite its importance, MIC determination is frequently subject to misinterpretation. This can be due to the lack of technical experience, improper performance of the standard protocol, or the reliance on visual inspection and reading of absorbance. As a result, MIC values may be inaccurately reported or even censored, leading to questionable conclusions about a compound’s efficacy. Another major challenge in MIC determination arises from the limited availability of the isolated or purified compounds. Often, the purification yield of a novel compound is relatively low, and because the MIC assay typically requires a minimum starting volume of 50 µl of a high-concentration stock, a significant amount of the compound can be consumed in a single experiment. This becomes a bottleneck when experiments are performed in triplicate, as is standard for reproducibility, thereby quickly depleting the already limited stock of the compound. The same concern applies to commercial antimicrobial agents, which are often expensive and sometimes impractical to use in larger volumes. Given these limitations, we have developed a method for determining the MIC in a more sensitive, miniaturized, and efficient way that requires lower volumes of the test compounds while still delivering accurate and reproducible MIC results. The current method has been developed considering all the standard parameters for determining MIC, such as bacterial strains, bacterial inoculum size of 0.5 McFarland standard turbidity, growth medium, incubation temperature, and two-fold serial dilution.

The widely accepted conventional methods for MIC determination, such as broth microdilution and agar dilution methods, can be labour-intensive, time-consuming, and may lack the throughput required for screening large libraries of compounds or for rapid diagnostic purposes as the amount of test sample is often limited in availability. Furthermore, these methods often rely on subjective visual interpretation of growth inhibition, which can introduce variability. In contrast, this five-step method provides foundational data for developing a novel antimicrobial strategy and exploring innovative approaches to determine the MIC for the purified compounds.

## Materials and Methodology

### Bacterial strain and Reagents

Bacterial strains *S. aureus* RN4220, *Vibrio fischeri* ATCC7744, *Klebsiella pneumoniae* MTCC109, and *Escherichia coli* NCIM2574 were used in this study. The culture medium for this bacterium, Tryptic Soya Agar (TSA), Tryptic Soya Broth (TSB), Muller-Hinton Agar (MHA), and Muller-Hinton Broth (MHB), was procured from Himedia. The Standard 90 mm sterile Petri plates from Tarsons were used to perform the assay.

### Procedure

#### Step 1: Preparation of the antibacterial stock solution (Timing:∼30-45 min)

1. An accurate 2x stock concentration of the test compound was prepared, which is necessary for the MIC. The concentration of the standard antibiotic should be prepared at 1 mg/mL in the desired solvent for the apparent solubility of the agent. The stock solution should be clear and transparent.
  a. A pre-calibrated analytical balance must be used for weighing.
  b. The HPLC-purified compound can be evaporated in a pre-weighed microcentrifuge tube, and the final weight can be taken to get the weight of the compound.
2. The compound was dissolved in a suitable soluble solvent (DMSO or water) based on the solubility by adding an appropriate solvent volume. Sterile distilled/MQ water should be used for water-soluble compounds. **Note:** Sometimes the compound does not dissolve completely; in that case, higher percentages of organic solvents should be used. In the conventional broth microdilution method of MIC, more than 10% DMSO also interferes with microbial growth. However, in this protocol, 50-100 % DMSO could be used to dissolve the compound and ensure the solvent does not interfere with the assay and putting a negative control.
3. The tube is vortexed thoroughly until completely dissolved. If needed, the tube should be kept in a water bath to apply gentle heat or sonicated to facilitate the dissolution of the compound.
4. The compound/antibiotic needs to be filter-sterilized using a 0.22 µm syringe filter.
5. The stock solution should be aliquoted into the sterile amber microcentrifuge tube for further use and the stock stored as aliquots at 4ºC. The stock and aliquots should be stored as per the stability of the compound at a particular temperature.

#### Step 2: Preparation of bacterial inoculum and culture (Timing:∼24-48 h)

1. The test microorganism is to be inoculated from the glycerol stock or lyophilized vial in TSB medium and incubated at 37ºC for 24 hours to obtain the growth of bacteria. 10-20 µL of bacterial culture is to be taken onto a TSA plate and incubated at 37ºC for 16-18 hours to obtain isolated colonies. With a sterile loop, 2-3 single colonies are to be picked and inoculated in 5-10 mL Muller-Hinton broth. The broth is to be incubated for 10-12 hours to obtain microbial growth.
2. The turbidity of the microbial suspension is to be measured using a spectrophotometer at 600 nm wavelength. The 0.5 McFarland is to be adjusted to standard turbidity by diluting the microbial suspension to obtain a CFU of 1 × 10^6^ CFU/mL for the MIC assay. **Note:** After adjusting the CFU, the prepared bacterial inoculum should be used within 30-60 minutes to maintain cell viability.

#### Step 3: 1:1 Serial dilution of the test compound (Timing:∼10-15 min)

1. The 200 µL or 1.5 mL sterile microcentrifuge tubes have to be labelled from 1-10 (for serial diluting the test compound).
2. 10-50 µL of the diluting solution DMSO/ water (based on the compound’s solubility) is added in each microcentrifuge tube.
3. An equal amount of the prepared 2x concentration compound is added from the stock solution to the first tube to maintain a ratio of 1:1 (Diluting solution: compound). **Note:** We recommend preparing the antibacterial compound stock fresh every time to avoid concentration variations caused by freezing and thawing.
4. A two-fold serial dilution is to be performed across the tubes labelled from 1-10, and appropriate volume is to be discarded from the final 10^th^ tube to maintain equal volumes.

**Note:** In case the sample amount is limited, a minimum of 10 µL of the diluting solution can be used; then an equal amount of the 2X or the highest concentration of the compound will be added to the first tube, and then the serial dilution needs to be performed. Pipetting is to be done gently several times to mix the compound properly and get the desired concentration.

#### Step 4: Preparation of bacterial lawn on MHA plates (Timing: ∼ 5-10 min)

1. The MHA plates should be labelled with sample ID, date, and compound tested, and also mark 1-10 spots on the inverted side of the agar plate.
2. A sterile cotton swab is to be dipped into the adjusted 0.5 McFarland standard turbidity suspension with 1 × 10^6^ CFU/mL.
3. The swab is gently pressed against the inside wall of the tube/flask to remove the excess amount of bacterial culture.
4. The swab is streaked evenly across the entire surface of the agar plate by rotating the plate at 60º three times to ensure complete coverage.
5. The round edges of the agar on the petri plate are to be swabbed to avoid patchy growth of the bacteria.
6. The plates are left to dry for 2-5 minutes inside the laminar air flow to ensure excess moisture is absorbed before applying the test compound.

#### Step 5: Application of the test compound/MIC assay

After making a uniform bacterial lawn, the agar diffusion method applies the test compound to the agar surface. As we have a minimal amount of compound, it saves our compound, and we can estimate the MIC range.

1. Using a 0.5-10 µL micropipette, 10 µL of the test compound prepared in step 3 is to be applied by serial dilution (1-10 microcentrifuge tubes) on the pre-labelled agar plate.
2. The dissolved compound is to be pipetted carefully onto the bacterial lawn without puncturing the agar.
3. The spots are to be placed at least 2 cm apart to avoid the overlapping inhibition zones. **Note:** The plate is not to be shaken or disturbed as the spots might overlap, resulting in false results.
4. The spots are allowed to air dry with the lid partially open in a sterile environment for 10-15 minutes before incubation.
5. The plates are placed in an inverted position in an incubator carefully to prevent condensation. **Note:** If the spots are not dry, plates can be positioned upright in the incubator for some time, and then inverted.
6. The plates are incubated at 35-37 °C for 16-18 hours for the fast-growing bacteria.
7. After incubation, the plates are observed for clear zone of inhibition and an inhibition with some clear area. **Note**: The MIC values or range are recorded by visual inspection. The MIC is the lowest antimicrobial compound concentration, with no visible growth of bacteria on the agar plate (the clear zone of inhibition at the place where the compound was placed). The MIC value is the consensus of the three technical replicates.

### Statistical analysis

MIC values calculated by both methods (this novel method and from broth microdilution assay) were statistically validated using the “Bland-Altman analysis” for method comparison. In this analysis, the MIC values were converted into log_2_ values, and bias was calculated. According to Bland-Altman statistics, a bias ≈ 0 indicates good agreement between both methods. On the other hand a bias ≥ ±1 would indicate that our proposed method systematically overestimates the conventional method, whereas a bias ≤ -1 would be indicative of our method underestimating the conventional method.

## Results and Discussion

The determination of MIC for antimicrobial agents continues to be regarded as one of the most crucial steps in identifying their therapeutic potential, and is the most important microbial susceptibility indicator [5, 6]. In this study, we tested Kanamycin (C = 500 µg/mL) against the bacterial strains *S. aureus, V. fischeri, K. pneumoniae*, and *E. coli* using our method and the broth microdilution method. Very often, purified compounds obtained from natural sources such as plant extract, fungi, bacteria and marine organisms, is limited by the yield after multiple purification steps. Given the limitations on the availability of the compound or drug, other application studies are more expensive and a significantly high amount is required to determine the MIC. The standard broth microdilution method was comparatively evaluated alongside our method, to determine the MICs of Kanamycin against the four bacterial strains and to compare MIC breakpoints in 96-well plates with bacterial inhibition on agar plates. In the broth microdilution assay for MIC determination, we used MTT as a viability indicator to visually detect metabolically active bacterial growth. The MTT kit (Roche) enabled distinct visualization of MIC endpoints, as the reduction of MTT to insoluble formazan produced an easily distinguishable colour gradient (Fig. 2A) [8, 9]. The results aligned with our method, using 5 µL of kanamycin dilutions on an agar plate. A larger zone of inhibition can be seen at position 1 with concentration ‘C’, while with the decreasing concentrations of the antibiotic, a smaller zone of inhibition could be visualized till the point it completely disappears. This point signifies that the last zone of inhibition that we can observe is the lowest concentration of the drug that inhibits the growth of bacteria (Fig. 2B). For both the methods, calculated log_2_ MIC values are given in the Table 1. The statistical bias, determined using the Bland-Altman analysis from the comparison of calculated MIC values, was found to be -0.00025 ± 0.81 (bias ± SD of bias).

**Table No.1:** Log_2_ MIC values calculated after determining the MIC from both methods.

| Bacteria | $\log_2$ MIC (Broth-dilution) | $\log_2$ MIC (Agar-diffusion) |
| --- | --- | --- |
| *S. aureus* | 4.966 | 3.966 |
| *V. fischeri* | 3.966 | 3.966 |
| *K. pneumoniae* | 4.966 | 5.965 |
| *E. coli* | 4.966 | 4.966 |

**Fig. 1:**
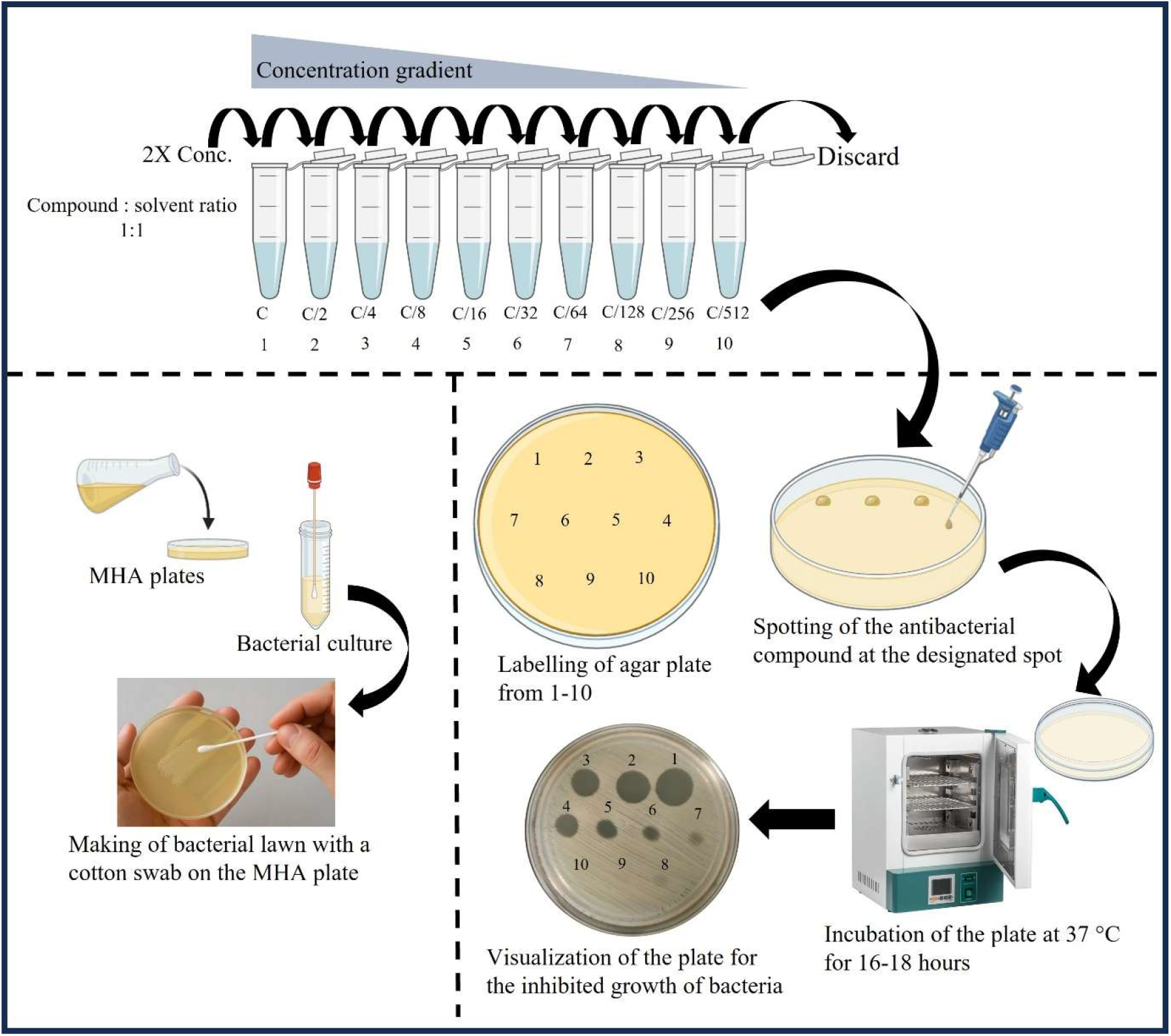
Visual representation of the defined MIC method and workflow.

**Fig. 2:**
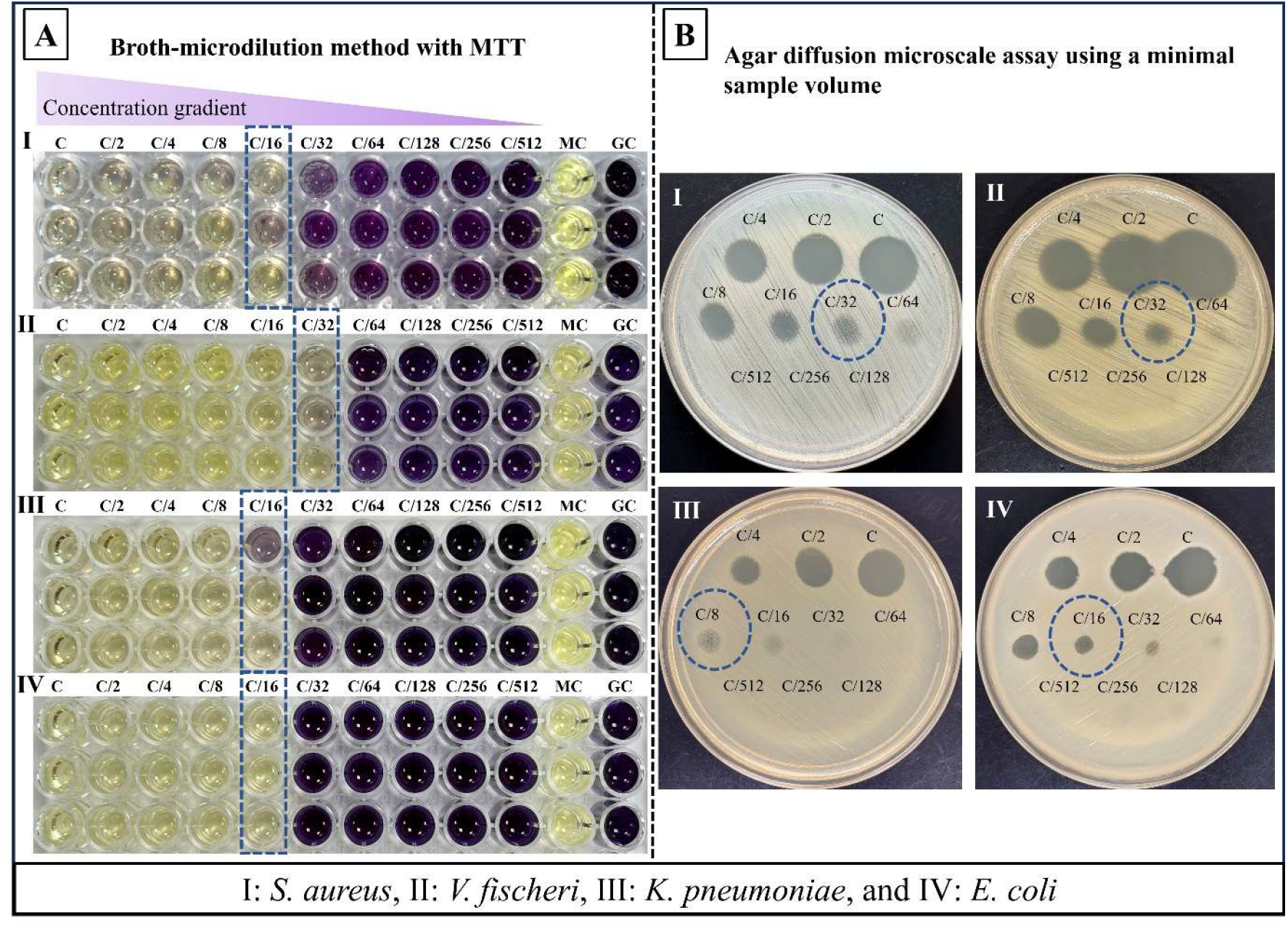
Side-by-side panel comparative images of both the methods for MIC determination.**A** represents the MIC of kanamycin antibiotic with the broth microdilution assay and visualization after the addition of MTT salt, and **B** shows the MIC of kanamycin antibiotic on an agar plate after a 2-fold serial dilution. Kanamycin concentration C = 500 µg/mL, MC = media control, GC = growth control.

Secondly, given the high cost and solubility issues of some commercial drugs or natural compound makes usage in larger amounts is unrealistic. Our method allows usage of lower volumes or concentrations, thereby circumventing the cost and solubility bottlenecks. The toxicity of the solvent in which the compounds are dissolved can also be assessed by diffusing it on agar alongside the test to confirm that the bacterial inhibition is due to the tested drug, not from the solvent.

We can therefore safely conclude that our method offers a rapid, reliable, reproducible, cost-effective, and visually distinct approach for MIC determination. This is particularly valid for those novel compounds, where the compound solubility and visualization pose significant challenges, while overcoming the key limitation of solvent toxicity, but maintaining strong agreement with established assays at the same time.

## Conclusion

This methodological study highlights the importance of sample volume in the determination of MIC while screening for purified and other commercially available compounds, which are often constrained by the available amount. Results obtained from the study support the method for determination of MIC. The defined method is easy to perform and cost-effective, particularly when the test compound is limited by the availability and volume.

## Acknowledgements

The authors acknowledge funding from the DBT BUILDER project no. BT/INF/22/SP42543/2021 for the work done in the manuscript. SK was supported by a fellowship from the same project.

## Author contributions

SK was instrumental in the collection of data, analysis of results and the primary scripting of the manuscript. SB was involved in the analysis of data and validation and the final draft of the manuscript.

## Data availability

The authors declare that the data supporting the findings of this study are available within the paper. Should any raw data files be needed in another format they are available from the corresponding author upon reasonable request.

